# High-throughput identification and marker development of perfect SSR for cultivated genus of passion fruit (*Passiflora edulis*)

**DOI:** 10.1101/322636

**Authors:** Yanyan Wu, Weihua Huang, Yongcai Huang, Jieyun Liu, Qinglan Tian, Xinghai Yang, Xiuzhong Xia, Haifei Mou

## Abstract

Simple sequence repeat (SSR) markers are characterized by high polymorphism, good reproducibility and co-dominance etc. They can be easily applied to develop efficient, simple and practical molecular markers. In the present study, bioinformatics methods were applied to identify high-throughput perfect SSRs of cultivar *Passiflora* genome. A total of 13104 perfect SSRs were obtained. SSR core sequence structure is mainly 2-4 bases, the maximum numbers are TA, AT, TC and AG. The maximum numbers of repetitions were up to 20 times. A total of 12934 pairs of SSR markers were developed by using bioinformatics software, and 20 pairs of markers were selected for amplification specificity assessment of MTX and WJ10, and the polymorphism rate was as high as 60%. The large-scale development of the SSR markers of *Passiflora* cultivar has paved a foundation for the efficient utilization of the germplasm resources of passion fruit, genetic improvement of the varieties and molecular breeding.

## Introduction

*Passiflora*, also known as passion fruit or egg fruit, is an important tropical and subtropical fruit tree. Cultivated species of *Passiflora* possesses high nutritional, medicinal and ornamental values. Thus, *Passiflora* is of importance and economic significance. There are about 520 species of passiflora in the world (Araya et al., 2017), while their morphological and agronomic traits etc. are more abundant, their genetic diversity is lower (Cerqueira-silva et al., 2014). In molecular biology studies, early detection of RAPD markers (Fajardo et al., 1998; Crochemore et al., 2003) identified the cultivar *Passiflora* with low DNA polymorphism. In the recent years, AFLP (Segura et al., 2002; Ortiz et al., 2012) and ISSR (Sousa et al., 2015; Santos et al., 2011)and other molecular markers have been utilized to detect DNA polymorphisms of *Passiflora*. Although these markers can detect DNA polymorphisms in *Passiflora*, RAPDs have poor reproducibility. The operation procedures of AFLP are complex and have high requirements for technic skills of the experimenters and for experimental equipment. Although ISSR is simple, its reaction conditions are difficult to grasp, and most of them are the explicit markers. SSR markers have the characteristics of high polymorphism, good repeatability, and co-dominance with low requirement for DNA detection. However, to date, the numbers of effective SSR primers developed and validated by researchers are still limited (Costa et al., 2017; Cazé et al., 2012; Cerqueira-silva et al., 2012; Araya et al., 2017; Martin et al., 2005; Padua et al., 2005), and these SSR primers are still not able to satisfy the requirement for genetic research and development of passionflower. Therefore, using sequencing data of *Passiflora* genome and bioinformatics methods to identify and develop more SSR markers is of importance, theoretical significance and application value for accelerating the research process of genetic diversity and marker-assisted selection breeding ofpassion fruit.

## 1 Materials and Methods

### 1.1 Plant materials

The genomic sequencing data of the cultivar *Passiflora* were uploaded from the Beltsville Agricultural Research Center to the NCBI Assembly: (https://www.ncbi.nlm.nih.gov/assembly/GCA_002156105.1/#/st;). *Passiflora* cultivars were planted in Germplasm Farm of Biotechnology Research Institute Guangxi Academy of Agricultural Sciences.

### 1.2 Extraction of DNA

*Passiflora* leaves were taken, cleaned with alcohol and stored in the −80°C refrigerator. The genomic DNA was extracted from *Passiflora* leaves according to the cetyl trimethyl ammonium bromide (CTAB) (Murray et al., 1980) method with appropriate simplification.

### 1.3 Bioinformatics Analysis Software

The main bioinformatics software SSR Search, developed and supplied by Beijing Novogene Technology Co., Ltd., was mainly used for the identification of SSRs; perl language script: extracting 100 bp for each flanking sequence of SSR; Primer 3: After being filtered, the SSRs were designed with primers, and one SSR-labeled primer design was performed each time.

## 2 Results

### 2.1 Identification of the genome-wide perfect SSR of cultivar passion fruit

We analyzed the 165.6 Mb data representing the genome of cultivar *Passiflora* and achieved an assembly level of Scaffold. The specific parameters were set as follows: (1) The minimum length of SSR repeat units was 2 bp; (2) The maximum length of the SSR repeat unit was 6 bp; (3) The minimum length of the SSR sequence was 12 bp; (4) The length of the SSR upstream and downstream sequences was 100bp. and (5) the minimum distance between the two SSRs was 12bp. Finally, we identified a total of 13104 SSRs and 2-6 core sequence numbers (Figure 1). The core sequences of most SSRs were 2-4 bases. The core sequences were mainly TA, AT, TC and AG. The highest number of repetitions was CT, which was 20 times; the least number of repetitions was TAT, which was 4 times (Figure 2).

**Figure 1.**
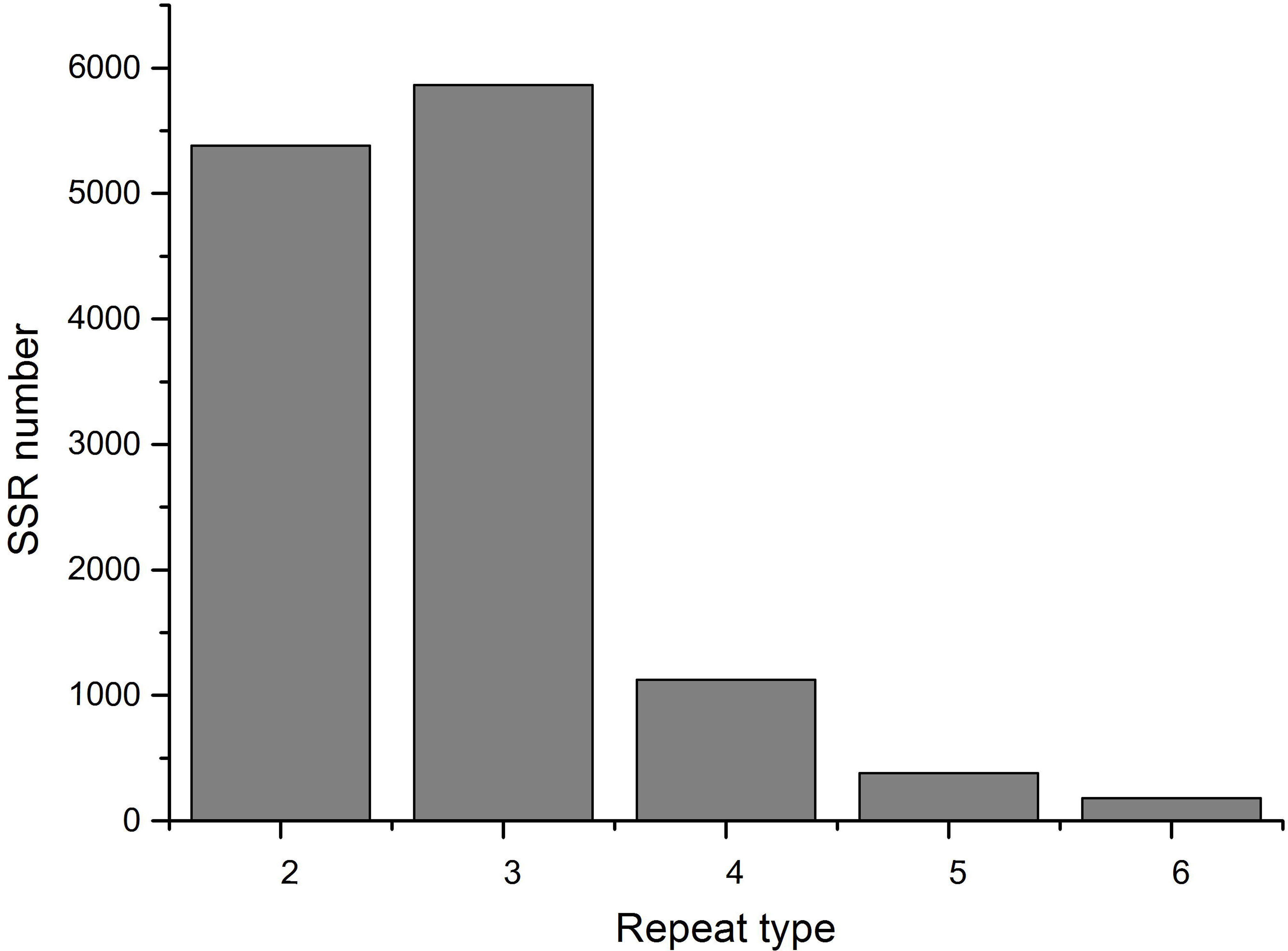
Complete SSR type and quantity statistics.

**Figure 2.**
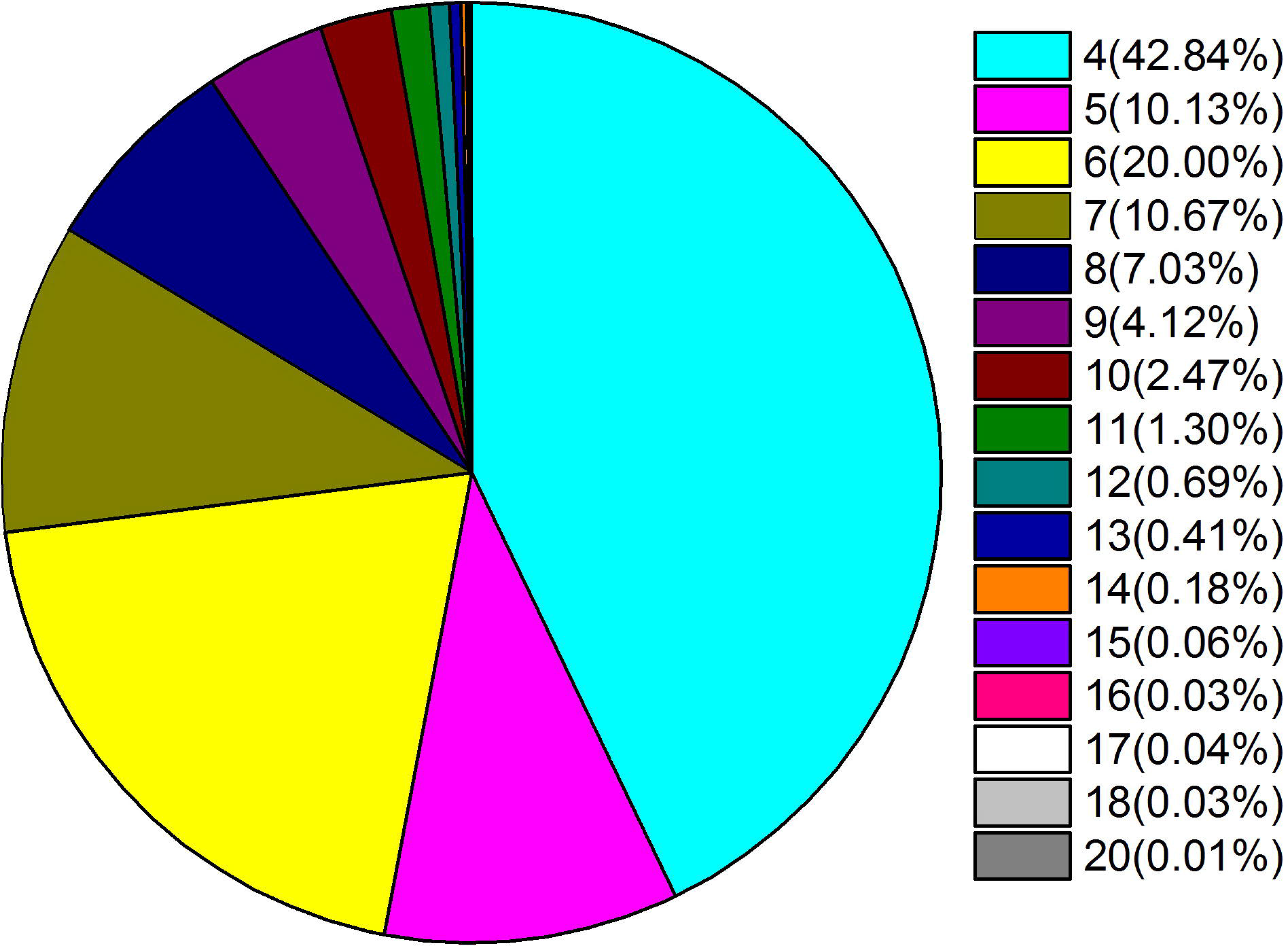
Total number of motif repeats for complete SSR.

### 2.2 Design of ‒genome-wide perfect SSR markers for cultivar genus *Passiflora*

Perl script ssr_filter.pl (in house) was used to filter the detected SSRs. The parameters were set to: -d 12 -len 100 means: (1) -d 12: The minimum distance between the two SSRs was 12 bp; (2) -len 500: The length of the upstream SSR primer was 500 bp between upstream and downstream. The identified 500bp flanking sequences from the SSR of the cultivar genome were extracted. Then, the primers were designed by using the primer design software Primer 3 and one SSR-labeled primer design were performed each time. The primer3 input file was used. Primers were designed based on following principles: (1) The optimal length of the primer was 24 bp; (2) The minimum length of the primer was 20 bp; (3) The longest primer length was 28 bp; (4) the optimal annealing temperature for primers was 55; (5) The lowest primer annealing temperature was 53; (6) The highest primer annealing temperature was 58 and (7) The maximum difference in the annealing temperature of a pair of primers was 1. Finally, we designed a total of 12934 pairs of SSR primers (supplementary **Table 1**).

### 2.3 The Development of genome-wide perfect SSR markers for vultivated genus *Passiflora*

We selected 20 pairs of SSR primers from 12934 pairs of primers (Table 1), and performed specific amplification to evaluate the DNA of Passion fruit and Passiflora chinensis, and found 12 pairs of SSR primers in the West. The polymorphisms in the DNA of the passionflower and Huangguo Passiflora (Figure 2), the polymorphism rate reached as high as 60%.

**Fig. 3.**
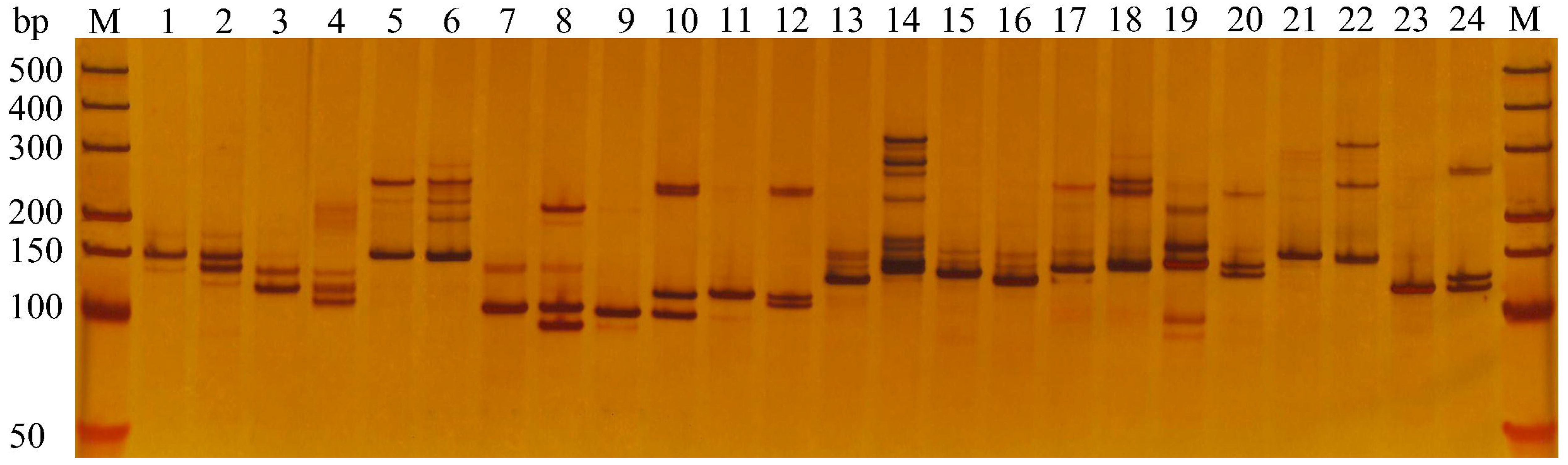
12 pairs of SSR markers in polymorphisms of genus *Passiflora* MTX and WJ10.

**Table 1.**
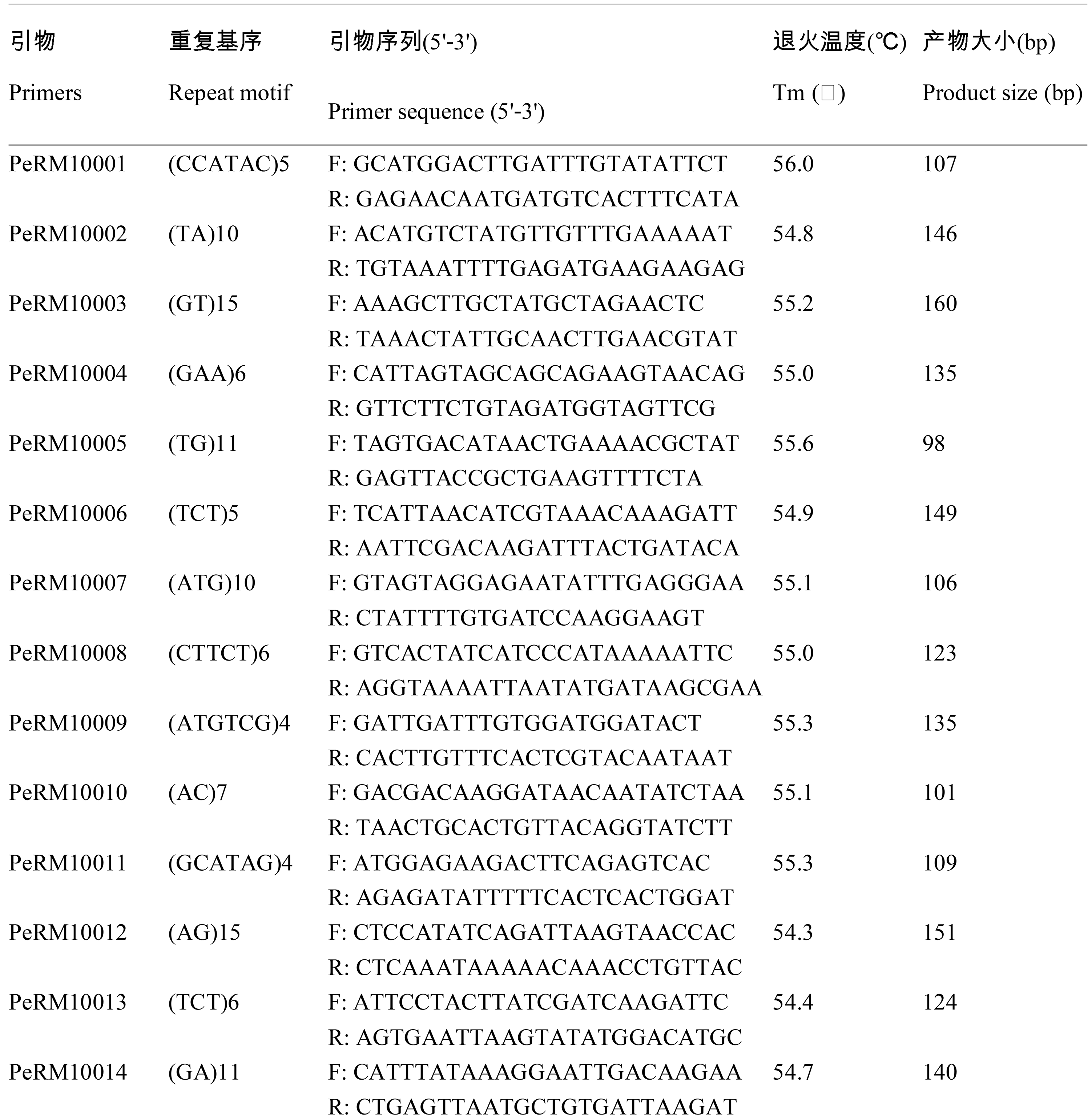
The 20 pairs of primers sequences for cultivated passion fruit

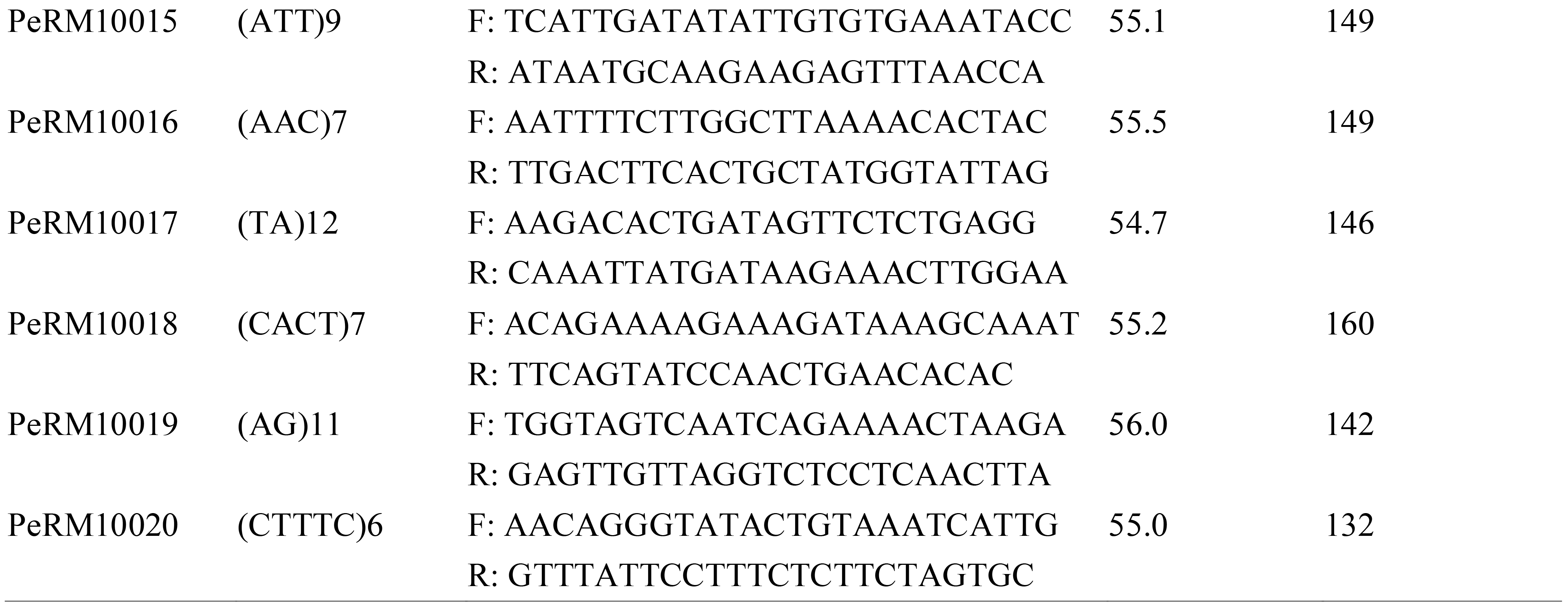

## 3 Conclusions

On May 22, 2017, scientists from Brazil and the United States jointly used Illumina GAII sequencing technology, for the first time, to perform whole genome sequencing of cultivar *Passiflora* CGPA1. The results were uploaded to NCBI, and raw data of 14.11Gb was obtained, and 165,656,733 bp cultivars were assembled. These genomic sequence information and results provide us with an opportunity for high-throughput identification of SSRs and for the development of SSR markers. This study was the first to use the sequencing results of the cultivar *Passiflora* genome. High-throughput identification of complete SSRs in the Passiflora genome and development of a large-scale cultivar S. Passiflora using SSR markers led to the establishment of an efficient cultivar Passionflower. The SSR marker system is a rich number of molecular markers for passion fruit, and provides a technical reserve for the construction of high-density genetic linkage maps of cultivar Passiflora and fine positioning of key peanut genes in the next step, paving the foundation for subsequent molecular breeding of passionflower basis.

## Reference

Araya S., Martins A.M., Junqueira N., Costa A.M., Faleiro F.G., Ferreira M.E.. 2017. Microsatellite marker development by partial sequencing of the sour passion fruit genome (*Passiflora edulis* Sims). BMC Genomics 18: 549

Cazé A.L., Kriedt R.A., Beheregaray L.B., Bonatto S.L., Freitas L.B.. 2012. Isolation and characterization of microsatellite markers for *Passiflora contracta*. Int J Mol Sci13: 11143–8

Cerqueira-silva C.B., Santos E.S., Souza A.M., Mori G.M., Oliveira E.J., Corrêa R.X., Souza A.P.. 2012. Development and characterization of microsatellite markers for the wild South American *Passiflora cincinnata* (Passifloraceae). Am J Bot 99: 170–2

Cerqueira-silva C.B., Santos E.S., Vieira J.G., Mori G.M., Jesus O.N., Corrêa R.X., Souza A.P.. 2014. New microsatellite markers for wild and commercial species of *Passiflora* (Passifloraceae) and cross-amplification. Appl Plant Sci 2: 1300061

Costa Z.D., Munhoz C.F., Vieira M.. 2017. Report on the development of putative functional SSR and SNP markers in passion fruits. BMC Res Notes 10: 445

Crochemore, Lúciamolinari M., Correavieira H.B., Esteves L.G.. 2003. Genetic diversity in passion fruit (*Passiflora* spp.) evaluated by RAPD markers. BrazArch Biol Techn 46: 521–7

Fajardo D., Angel F., Grum M., Tohme J., Lobo M.. 1998. Genetic variation analysis of the genus *Passiflora* L. using RAPD markers. Euphytica 101: 341–7

Martin P., Makepeace K., Hill S.A., Hood D.W., Moxon E.. 2005. Microsatellite instability regulates transcription factor binding and gene expression. Proc Natl Acad Sct USA 102: 3800–4

Murray M.G., Thompson W.F.. 1980. Rapid isolation of high molecular weight plant DNA. Nucleic Acids Res8: 432–5

Ortiz D.C., Bohórquez A., Duque M.C., Tohme J., Cuéllar D.. 2012. Evaluating purple passion fruit (*Passiflora edulis* Sims f. *edulis*) genetic variability in individuals from commercial plantations in Colombia. Genetc Resour Crop Ev59: 1089–99

Padua J.G., Oliveira E.J., Zucchi M.I., Gcx O., Lea C.. 2005. Isolation and characterization of microsatellite markers from the sweet passion fruit (*Passiflora alata* Curtis: Passifloraceae). Mol Ecol Notes 5: 863–865

Santos L., Oliveira E., Silva A., Carvalho F., Costa J.L.. 2011. ISSR Markers as a tool for the assessment of genetic diversity in *Passiflora*. Biochem Genet 49: 540–4

Segura S., D’eeckenbrugge G.C., Bohorquez A., Ollitrault P., Tohme J.. 2002. An AFLP diversity study of the genus Passiflora focusing on subgenus *Tacsonia*. Genet Resour Crop Ev 49: 111–23

Sousa A.G., Souza M.M., Melo C.A., Sodré G.A.. 2015. ISSR markers in wild species of *Passiflora* L. (Passifloraceae) as a tool for taxon selection in ornamental breeding. Genet Mol Res 14: 18534–45

